# Numerical response of predators to large variations of grassland vole abundance, long-term community change and prey switches

**DOI:** 10.1101/2020.03.25.007633

**Authors:** Patrick Giraudoux, Aurélien Levret, Eve Afonso, Michael Coeurdassier, Geoffroy Couval

**Affiliations:** Chrono-environnement, Université de Bourgogne Franche-Comté/CNRS usc INRA, 25030, Besançon Cedex, France; FREDON Bourgogne Franche-Comté, 12 rue de Franche-Comté, 25480, Ecole-Valentin, France

## Abstract

Voles can reach high densities with multi-annual population fluctuations of large amplitude, and they are at the base of large and rich communities of predators in temperate and arctic food webs. This status places them at the heart of management conflicts wherein crop protection and health concerns are often raised against conservation issues. Here, a 20-year survey describes the effects of large variations in grassland vole populations on the densities and the daily theoretical food intakes (TFI) of vole predators based on roadside counts. Our results show how the predator community responds to prey variations of large amplitude and how it reorganized with the increase in a dominant predator, here the red fox, which likely negatively impacted hare, European wildcat and domestic cat populations. They also indicate which subset of predator species might have a role in vole population control in the critical phase of a low density of grassland voles. Our study provides empirical support for more timely and better focused actions in wildlife management and vole population control, and it supports an evidence-based and constructive dialogue about management targets and options between all stakeholders of such socio-ecosystems.

## Introduction

The relationship between people and rodents is an old one. Early accounts clearly show that rodents were a destructive agent for crops and a source of disease for many ancient and current societies [1–3]. Voles can reach high densities with multi-annual population fluctuations of large amplitude, and they are often considered as pests in temperate farmland [4, 5]. However persecuted for this reason [4, 6], their effects on biodiversity are crucial. They are at the base of temperate and arctic food webs, maintaining large and rich communities of predators, as well as modifying nutrient cycling, soil aeration, and micro-organism assemblages [7]. This status places them at the heart of management conflicts where crop protection and health concerns are often raised against conservation issues [6]. Moreover poisoning when using chemicals for rodent pest control can depress populations of predators that are able to contribute to the regulation of rodent populations [8, 9]. A better understanding of the links between grassland vole population variations and predator responses will allow more timely and better focused management actions for all stakeholders in multifunctional socio-ecosystems.

Predation has been suggested to be one of the main drivers of rodent population fluctuations. Theory predicts that specialist predators that feed on one or a few kinds of prey can destabilize prey populations because they exert delayed- and direct density dependent mortality on their prey populations, while generalist predators, which feed on a wide variety of prey species, have direct density dependent mortality and therefore stabilize prey populations [10]. However, experimental tests on this prediction, e.g. predator removal and comparative field studies, have provided evidence both supporting and rejecting this hypothesis [11–17].

Studies in the Arctic and Fennoscandia on small mammal population cycles have accumulated support for the predation hypothesis [18]. For instance, in the tundra of Greenland, the numerical response of the stoat (*Mustela erminea*) has been shown to drive the population dynamics of the collared lemming (*Dicrostonyx groenlandicus*) by a 1-year delay. These dynamics are concurrently stabilized by strongly density-dependent predation of 3 generalists, the arctic fox (*Vulpes lagopus*), the snowy owl (*Bubo scandiacus*), and the long-tailed skua (*Stercorarius longicaudus*) [15, 19].

Population dynamic patterns of the common vole (*Microtus arvalis*) in intensive agricultural landscapes of south-west France are largely consistent with five of six patterns that characterize rodent cycles in Fennoscandia and can be explained by the predation hypothesis [20]. Hence, there is little doubt that in European arctic and temperate ecosystems predation in combination with other factors, can play a role in regulating small mammal population dynamics [17]. However, in temperate ecosystems the multiplicity of prey-resources and the larger number of predator species combined with landscape diversity (e.g. the spatial arrangements of optimal and suboptimal habitats for prey and predators) [21, 22] make the disentangling of the detailed processes and the role of each species and factors involved a challenge [1, 17].

For instance, based on a 20-year survey of the effects of an epidemic of sarcoptic mange that decreased fox populations in Scandinavia, Lindström *et al.* [23] *revealed that red fox (Vulpes vulpes*) predation was a crucial factor in conveying the 3-4 year fluctuations of voles (both bank and field voles (*Myodes glareolus* and *Microtus agrestis*)) to small game, e.g. periodically limiting the populations of hare (*Lepus europeus*), tetraonids (*Tetrao urogallus, Tetrao tetrix, Bonasia bonasia*) and rowdeer fawns (*Capreolus capreolus*). The importance of such prey switchings on prey population dynamics has also been reported for a long period in Newfoundland, where lynx (*Lynx lynx*), prey on snowshoe hares (*Lepus americanus*), until the hare population crashes. Then, lynx switch to caribou calves (*Rangifer tarandus*), and the cycle continues [24]. As a whole, those multiple and complex interactions can hardly be investigated in depth by simple modelling [25] or by small-scale experiments that cannot technically take into account all the relevant space-time scales and species communities involved in the real world and, thus, be generalized.

However, stakeholders in such systems are often protagonists of endless debates about regulation adoption and management decisions, which each of them advocating the control of one among many possible population targets and subsequent options for management. This debating is the case in the Jura mountains where massive outbreaks of a grassland vole species, the montane water vole, occurs with 5-6 year cycles and population densities exceeding 500-1000 ind.ha^−1^. High density peaks propagate over grasslands under the form of a travelling wave [5, 26]. In the same area, outbreaks of the common vole (> 1000 ind.ha^−1^), another grassland vole, also occur, however, they are non-cyclic in this area [7]. A number of field studies and studies using modelling have shown that the population dynamics of the two species are shaped by landscape features, with hedgerow networks and wood patches dampening the population dynamics and by contrast open grassland landscapes amplifying the outbreaks [5, 27–32]. Those outbreaks provide regularly massive quantities (up to > 80kg.ha^−1^) of prey for many species of carnivorous mammals and birds in grassland and by contrast low densities of secondary prey-resources that are less accessible (vegetation and/or anti-predation behaviour) such as forest, marsh and fallow small mammals (maximum about 3kg.ha^−1^) (e.g. bank vole, wood mice, (*Apodemus* sp.), field vole, etc.), with periodic (5-6 years) concomitant low densities in every habitats.

The variation in this predator community structure over the time span of large fluctuations of prey abundance has not been documented yet in this system, limiting both comparisons with ecosystems described in other part of the world where small mammal outbreaks occur [4] or with more simple food webs of northern ecosystems. Moreover, a large scale inadvertent experiment was offered by chemical control of vole populations in the 1990s, leading to a dramatic decrease in the fox population due to indirect poisoning, and its gradual recovery the following years after a shift in vole control practices [8].

The aim of this 20-year study is to describe the effects of large variations of grassland vole populations on their predator communities and of the long term increase in the fox population in such system. The aims were to (i) describe how a predator community responds to prey variations of large amplitude, (ii) describe how this community reorganizes over the long term with increases in a dominant predator, here the red fox, (iii) attempt to quantify the prey consumption of this predator community. The possible impact on the grassland prey population dynamics will be discussed on this basis.

## Material and methods

### Study area

The study was carried out around the Pissenavache hamlet (46.95°N, 6.29°E) in Franche-Comté, France, in an area of 3425 ha (2646 ha of farmland, 1094 ha of forest, 167 ha of buildings), at an average altitude of 850-900 m above sea level (Fig. 1 and 2). There, 100% of the farmland was permanent grassland used for pasture and (high grass) meadow for cattle feeding in winter (minimum of 5 months, November-March), with a productivity ranging from 5-6.5 tonnes of dry matter.ha^−1^.an^−1^ under the specifications of the European Protected Geographical Indication of the locally produced Comté cheese. A KML file (S1 kml file) with the bounding box of the study area is provided with the data.

**Fig 1.**
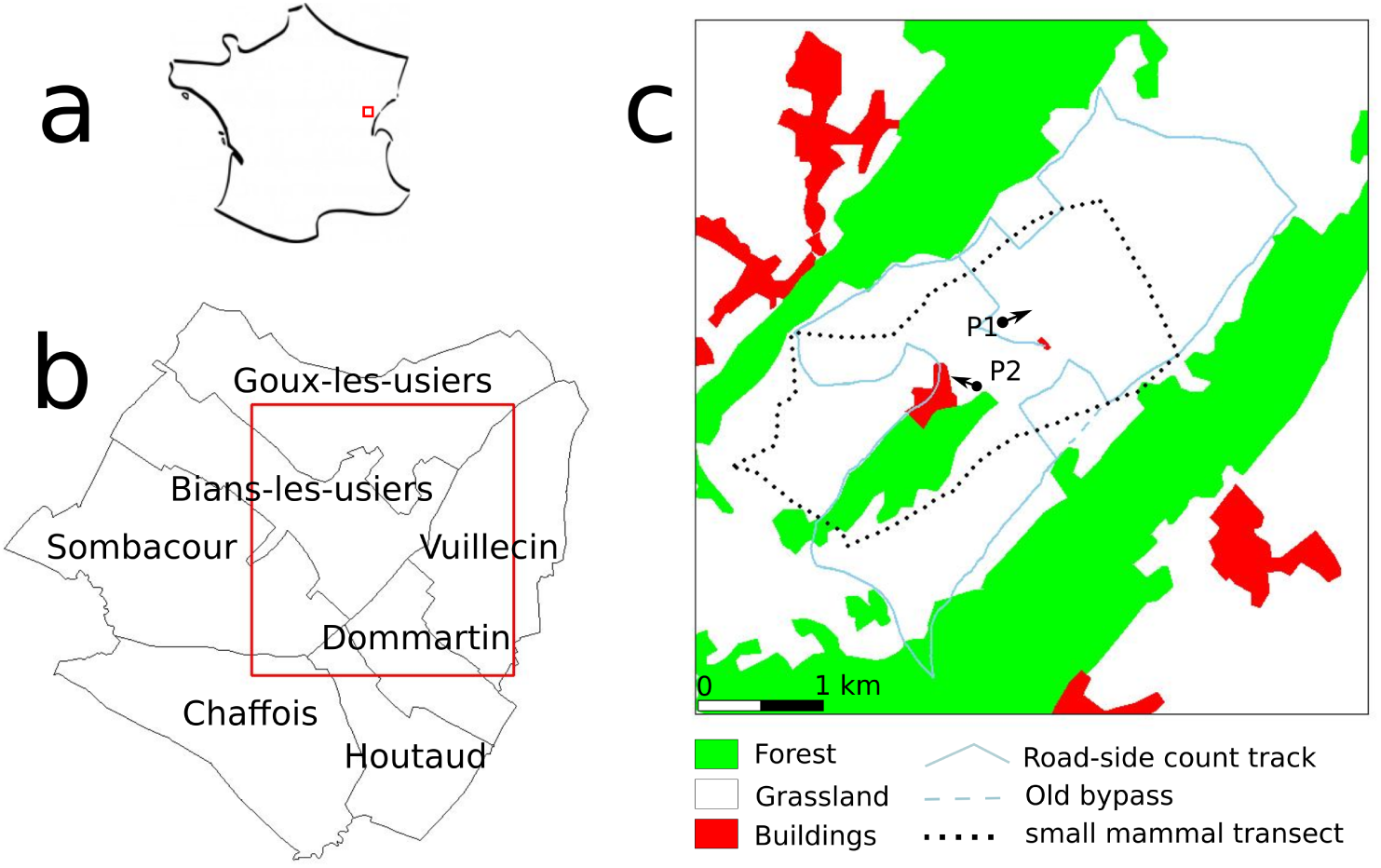
Location of the study area. a, general location in France; b, study area (red square) and communes it includes; c, land cover, road side counts and the small mammal transect, P1 and P2 indicate the directions of Fig. 2 photos. Until 2009, a road side count segment was driven straight along the dotted line, but in 2010 mud prevented the use of this bypass and slightly changed the itinerary (n-shaped solid line around the dotted line). Commune boundaries were derived from OpenStreetMap and land use from ‘BD Carto’ provided freely for research by the *Institut Géographique National*, modified based on field observations.

**Fig 2.**
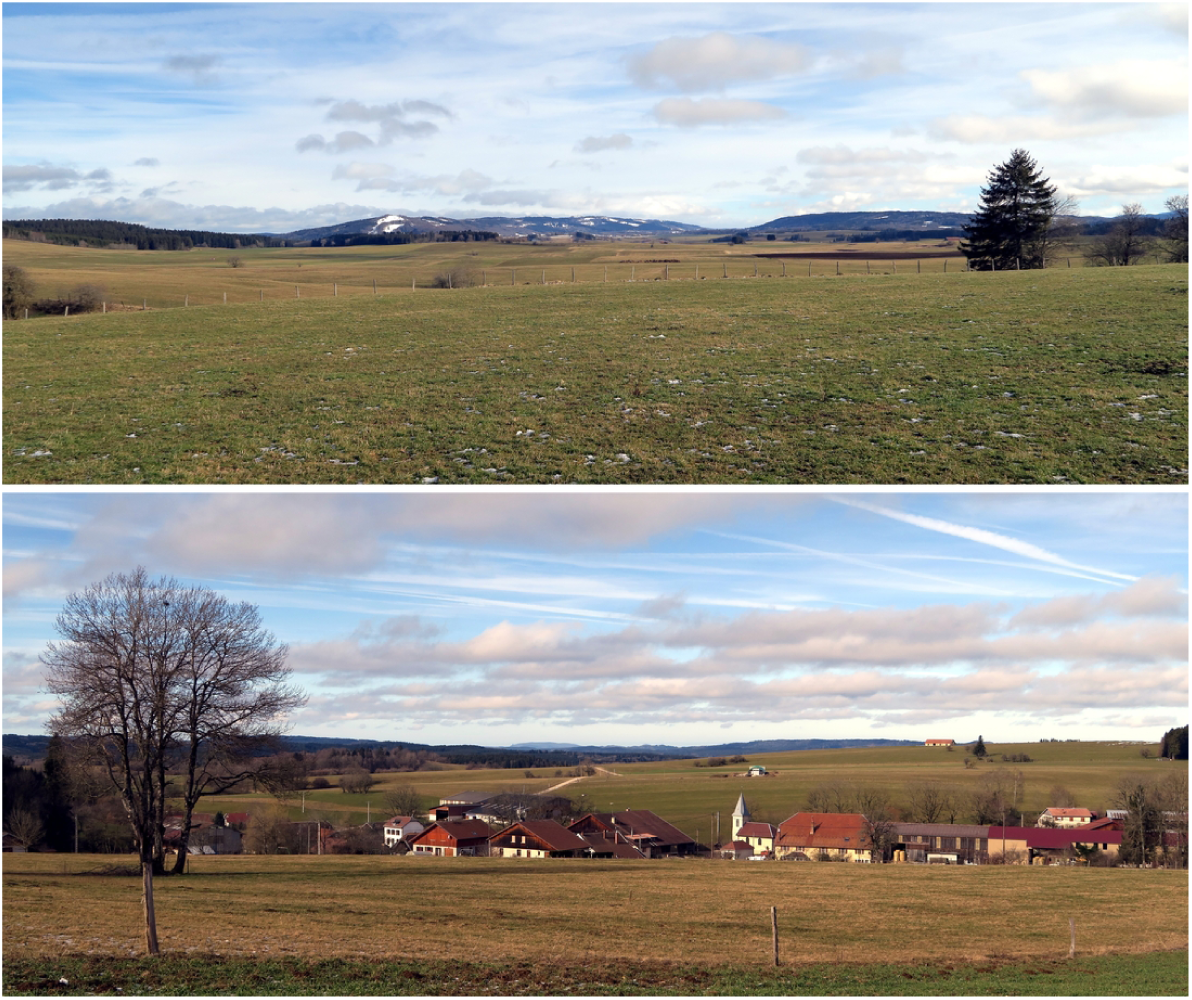
General views of the study area. Top, from the road-side count road at P1 (see Fig. 1); bottom, from P2 with the Pissenavache hamlet, a segment of the road-side count road can be seen in the background (photos PG, 20/02/2020).

### Roadside counts

Predator and hare (*Lepus europeus*) populations have been monitored from June 1999 to September 2018 (20 years) using night and day road-side counts. Each sampling event consisted of driving a car with 4 people (the driver, a data recorder and two observers) along a fixed track at less than 20 km/h. The length of the track was 18.6 km from 1999 to 2009 and then 19.6 km due to a slight variation in the itinerary (trail blocked by mud, see Fig. 1). Observations were performed using 100-W spotlights at night and binoculars for species identification. Distinction between domestic cats (*Felis silvestris catus*) and European wildcats (*Felis silvestris silvestris*) were made visually considering phenotypic criteria (relative to pelage and morphology) without possible distinction of hybrid individuals. Double counting was unlikely because the trail was straight, visibility large (Fig. 2) and observers were careful about animal movements. Sampling was carried out on 3 successive nights after sunset (4 when meteorological conditions prevented sampling) called a ‘session’. The same track was also driven by daylight on another day in the early morning. Most often 3-4 sessions a year were carried out corresponding to seasons, but only 1 session in autumn since 2016. Day road-side counts were stopped in 2017. Each observation was recorded on a paper map (IGN 1/25000). A Kilometric Abundance Index (KAI) was calculated for each session as the maximum number of animals recorded km^−1^ (thus providing a lower limit for the number of animals present). For the 2001-2006 period, only the total counts without the localizations of observations were available. Thus, only the 1999-2000 and 2007-2018 observations could be georeferenced.

### Daily food intake

Theoretical daily food intakes (TFI) per predator species were computed following Crocker *et al.*’s method [33] with small mammals considered as prey. The average body mass of predators, when missing in [33], was estimated based on the *Encyclopédie des carnivores de France* [34–37], the Handbook of Birds of Europe, the Middle East and North Africa [38] and the Encyclopedia of Life (https://eol.org).

### Small mammal relative densities

#### Transects

Small mammal *(A. terrestris, M. arvalis* and *Talpa europea*) relative abundance was assessed using a transect method adapted from [39–41]; a 5 m-wide transect across the study area was divided into 10-m-long intervals and the proportion of intervals positive for fresh indices (tumuli, molehill, runway, faeces, cut grass in holes) was considered an index of abundance. The total transect length was 11.6 km (Fig. 1). Sampling was carried out once a year in April 2007 and then in August from 2008 to 2010, followed by at least twice a year generally in spring and autumn from 2011 to 2018.

#### *A. terrestris* communal scores

To obtain abundance assessments on a larger space-time scale, abundance was also assessed at the commune-scale by technicians of the FREDON of Bourgogne Franche-Comté (a technical organization for plant pest prevention and control contracted by the Ministry of Agriculture [42]), in the 7 communes crossed by the road-side count itinerary (Fig. 1). Assessments were made in autumn since 1989. The FREDON assessment uses a ranking system that ranges from 0 to 5: 0 - no *A. terrestris* sign in any parcel within the commune; 1 - low or no *A. terrestris* tumuli, voles and moles (*T. europea*) cohabiting the same tunnel systems; 2 - *A. terrestris* tumuli present in some parcels within the commune and mole burrow systems still present in some parcels; 3 -*A. terrestris* tumuli present in some parcels within the commune, few or no mole burrow systems present in the commune; 4 - *A. terrestris* colonies established in the majority of meadows and within pastures; 5 - all of the commune colonized by *A. terrestris*. The FREDON index not directly translates to transect-based indices, partly because it is applied at the commune scale and not the parcel scale, but Giraudoux et al. [41] found that levels 0-1 correspond to densities < 100 voles.ha^−1^, level 2 to 100-200 voles.ha^−1^, and levels 3-5 to > 200 voles.ha^−1^. For a given year, the median score of the 7 communes was taken as a score of abundance.

#### Grassland prey resource relative abundance

The dynamics of prey resource abundance in grassland have been estimated (i) over the time span when transects were carried out, summing the relative abundance of *A. terrestris* and *M. arvalis* divided by four, divided by the maximum of this sum over the series and (ii) before this time span, when no transect was present, by dividing the FREDON score by the highest score recorded during the study (5). This process took into account that the *M. arvalis* body mass is four times smaller than *A. terrestris*’s on average [43] and helped to better visualize grassland rodent populations variation on the same scale and fill the gap when transect data were lacking. The amplitude of the high density phase is biased to an unknown extent with this method (e.g. arbitrarily summing weighted relative abundances, chained with standardized FREDON scores), but not the time-locations of the low density phases. Thus, the alternation between high density and low density phases, which are always very large (ranging from 0-1000 voles.ha^−1^), was robustly and correctly represented over the time series as an abundance index, in the best possible way given the data, for further comparisons.

### Rodenticide use

In France, bromadiolone, an anticoagulant rodenticide, has been used to control water vole populations since the 1980s, with deleterious effects on non-target wildlife including vole predators [9]. In the early 2000s, the development of an integrated pest management (IPM) approach [44] led to a dramatic decreasein the quantity of bromadiolone applied by farmers and their non-intentional effects [8, 9]. By law, the delivery of bromadiolone baits for vole control to farmers is under strict FREDON supervision and compulsory usage declaration to ensure traceability [45]. Data on bromadiolone quantities used in the 7 communes of the study area were provided by the FREDON of Bourgogne Franche-Comté.

### Statistical analyses

Statistical and spatial analyses were performed in R (version 3.6.2) [46] with the packages Distance [47], pgirmess [48], rgdal [49], rgeos [50], using QGIS 3.10 [51] complementarily.

#### Grassland small mammal abundance

The standard errors of small mammal relative abundances assessed from transects were computed across 1000 bootstrap replicates [52]. The grassland prey resource index corresponding to each road-side count was linearly interpolated over time between the two bracketing abundance index estimates.

#### Response of predators to prey abundance

We used generalized linear models with a Poisson error distribution of the form *n* = *a*_0_ + *a*_1_*ln*(*x*_1_) + *a*_2_*x*_2_ + *a*_3_*x*_3_ + *1:*, with *n*, the number of observations, *x*_1_, the length of the itinerary, *x*_2_, the season, *x*_3_, the prey abundance index, *a*_*i*_, the model coefficients, and *1:*, the residuals. To avoid overestimation of the degrees of freedom from time series data (here irregular and intrinsically autocorrelated), statistical inference was computed using permutation tests.

#### Predator and hare spatial distribution

The shortest distance of observations to the road-side count itinerary, to the nearest forest and to the nearest building were computed [49, 50] and their distribution examined.To test whether the proximity of some habitats might explain the observed distributions and their variations, the mean distance to forest and buildings were compared to the mean distances obtained from 1000 simulations of the same number of random positions as the number of observations in the strip observed along the itinerary.

#### Predator and hare population density estimates

To obtain density estimates, the distance to the itinerary data were analysed using conventional distance sampling with a truncation distance [53–55] including 90% of the observations for each species at the minimum. As avoidance behaviour along the road was detected for most species, we used hazard-rate detection functions fitted to the data. This function type has a more pronounced shoulder that compensates for the bias due to avoidance [47]. Models with a seasonal effect as a covariate were compared with concurrent models with no covariate using the Akaike Index Criterion [56].

## Results

### Small mammal density and prey resource variation

Fig. 3a shows the cyclic variations of *A. terrestris* from 1989 to 2018. Predators communities have been monitored during the last four cycles, but the local populations dynamics of small mammals during the last three cycles only (Fig. 3b). A clear synchrony of the low density phase between rodent species was observed, while *T. europea* and *A. terrestris* peaks were in phase opposition. In terms of prey resource, low density phases contrast with the phases of large abundance of grassland voles (Fig. 3c).

**Fig 3.**
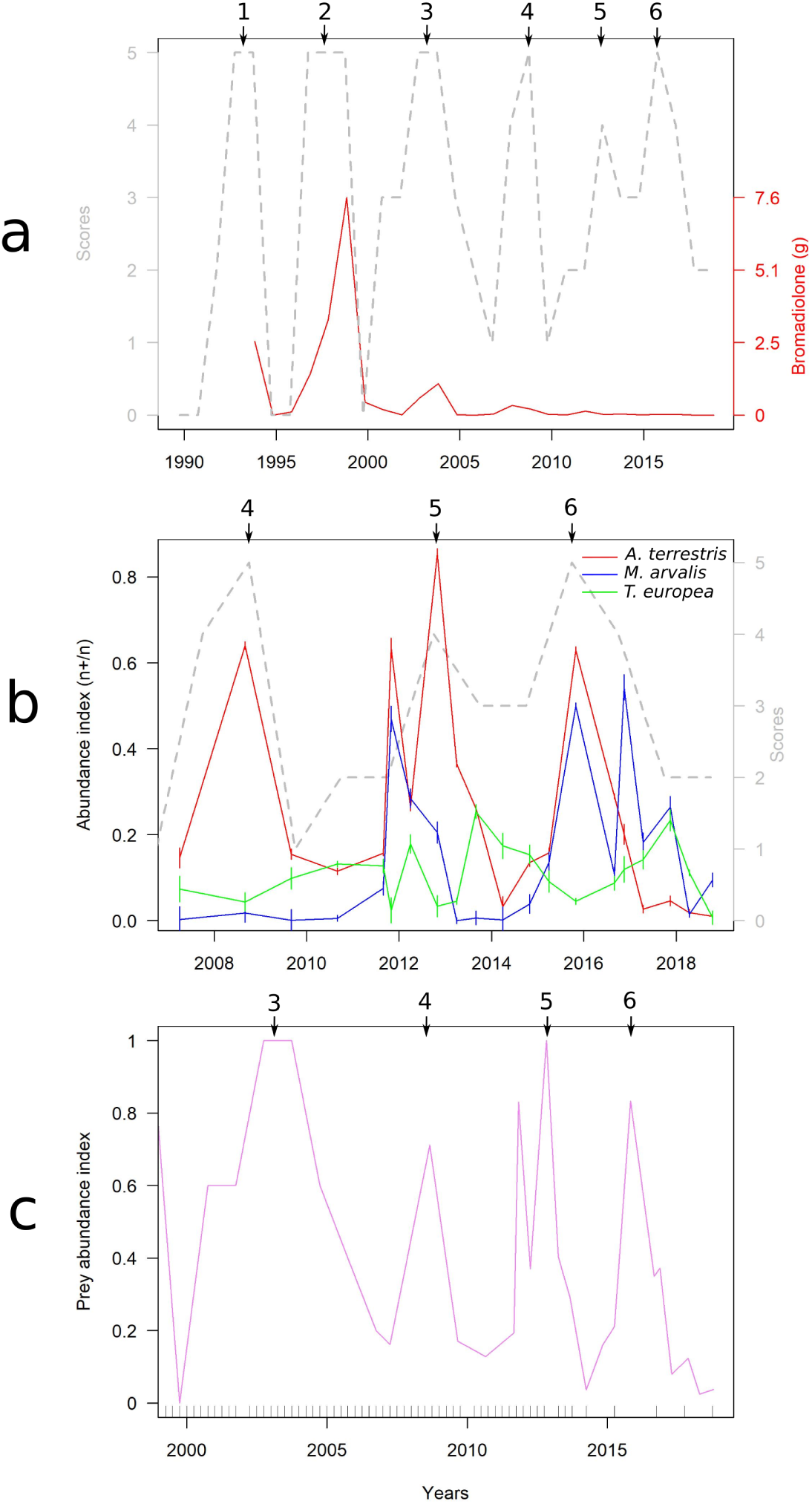
Small mammal population dynamics. Numbers with arrows indicate high density peaks in the communes including the study area; a, dotted grey line, *A. terrestris* FREDON scores; red line and red scale, quantity of bromadiolone (g) applied for *A. terrestris* control in the communes of the study; b, abundance index based on transects, vertical bars are 95% confidence intervals (grey scale and dotted line are related to the *A. terrestris* FREDON scores for comparison); c, estimated variations of the grassland prey resource, the rug on the x axis represents roadside count events.

### Numerical response of predators to grassland prey variation and hare relative abundance

#### Time variations

Twenty-seven species for the day road side counts and 24 for the night were observed, corresponding to 19,010 and 7,355 individual observations respectively, and to 58 sessions for each count type (≃ 348 night or day counts in total). Among them, the following species were both observed frequently enough over time and considered of interest for this study: for day roadside counts, the carrion crow (*Corvus corone*), the common buzzard (*Buteo buteo*), the red kite (*Milvus milvus*), the kestrel (*Falco tinnunculus*), the domestic cat (*Felis silvestris catus*), the hen harrier (*Circus cyaneus*); for night roadside counts, the European hare (*Lepus europeus*), the red fox (*Vulpes vulpes*), the domestic cat (*Felis silvestris catus*), the European wildcat (*Felis silvestris silvestris*), the long-eared owl (*Asio otus*), the European badger (*Meles meles*). Some were occasional visitors and likely play a marginal role on vole prey (e.g. grey herons (*Ardea cinerea*) could regularly be observed preying on voles in grassland). Others, such as some mustelids (stoat (*Mustela erminea*), least weasel (*M. nivalis*), stone marten (*Martes foina*), pine marten (*M. martes*)) were elusive and hardly detected by roadside counts.

Fig. 4 shows the dynamics of diurnal species. For each species KAI differences between seasons were found statistically significant except the domestic cat (Table 1 and Fig. 5).

**Table 1.** Statistical significance (p(H_0_)) of the model coefficients obtained by permutations, and model r-squared. Numbers between parentheses are values when one outlier is dropped (see results). n.s., not significant.

| count type | species | season | prey abundance | $r^2$ |
| --- | --- | --- | --- | --- |
| day | carrion crow | < 0.001 | n.s. | 0.68 |
|  | common buzzard | 0.002 | < 0.001 | 0.59 |
|  | red kite | 0.001 | 0.05 | 0.44 |
|  | kestrel | < 0.001 | 0.09 (0.01) | 0.62 (0.66) |
|  | domestic cat | n.s. | n.s. | - |
|  | hen harrier | 0.009 | 0.08 (0.02) | 0.52 (0.51) |
| night | hare | n.s. | 0.007 | 0.42 |
|  | red fox | n.s. | n.s. | - |
|  | domestic cat | < 0.001 | n.s. | 0.54 |
|  | wildcat | n.s. | < 0.001 | 0.42 |
|  | long-eared owl | n.s. | n.s. | - |
|  | European badger | n.s. | n.s. | - |

**Fig 4.**
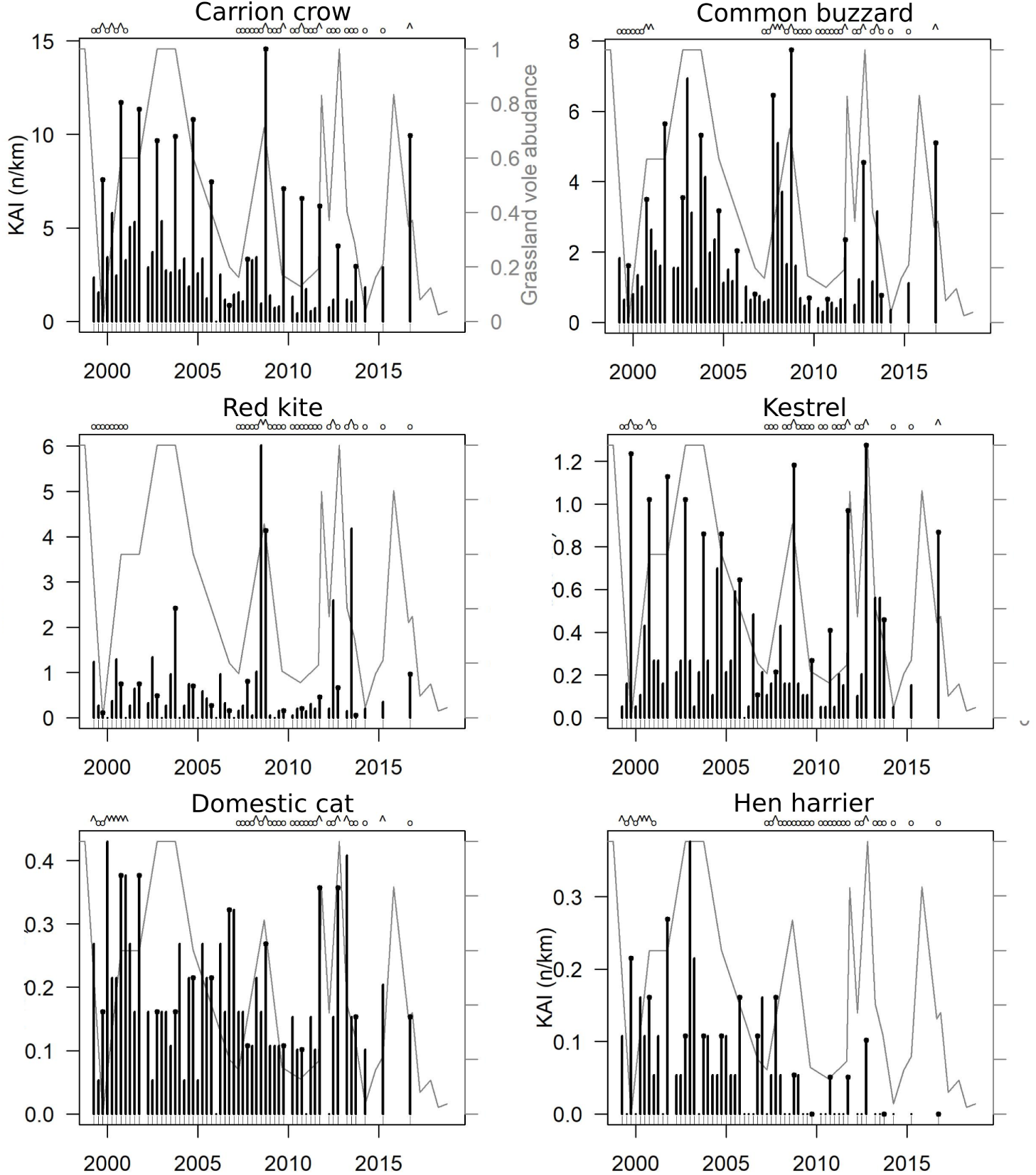
Day roadside counts. Black circles at the bar top identify autumn counts. The grey line in the background shows the variations of grassland prey abundance (the scale is the same in every plot). The letters above identify the sessions available and selected to estimate densities based on distance sampling during high (^) or low (o) abundance period.

**Fig 5.**
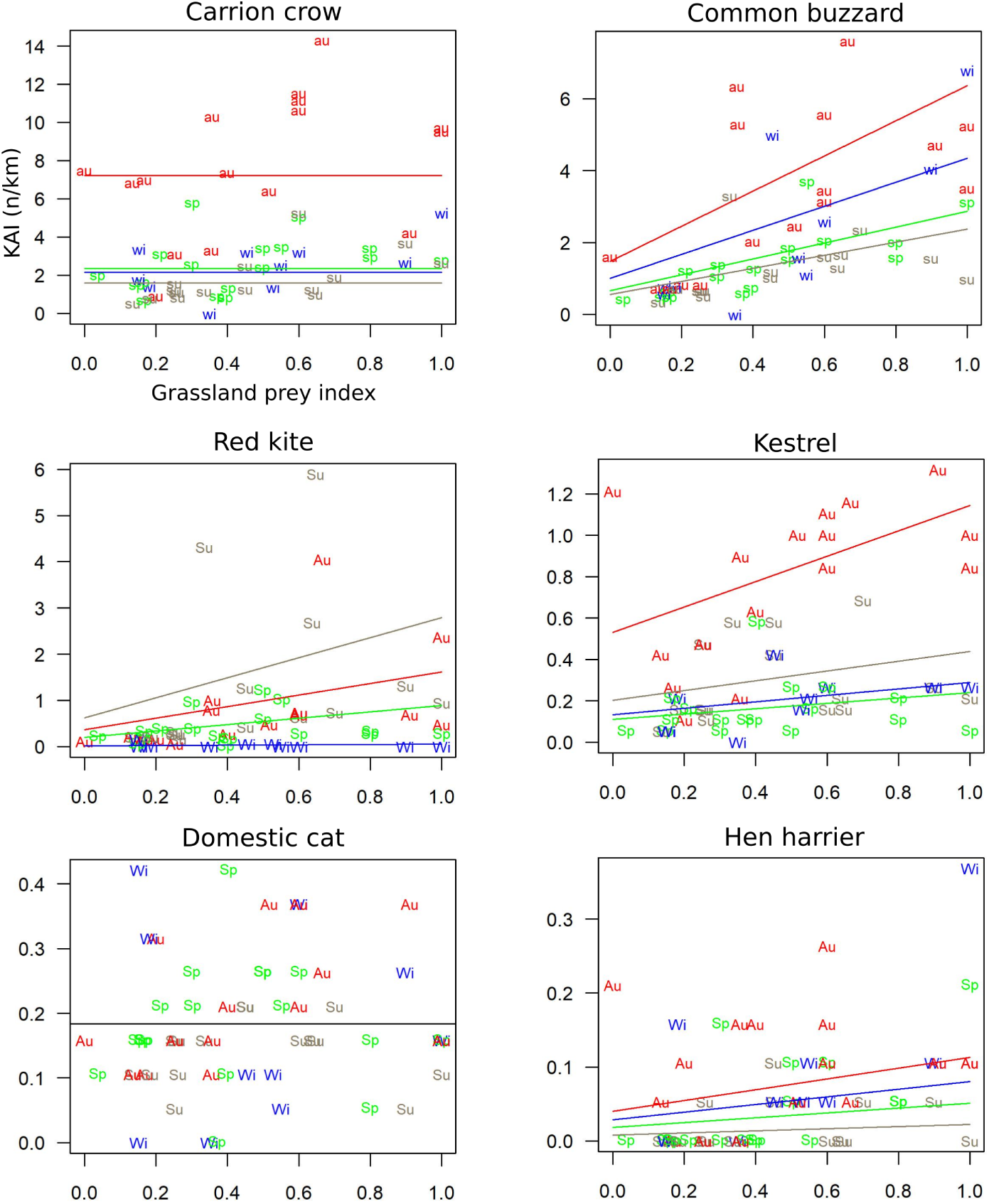
Biplots of diurnal KAIs as a function of grassland prey index. sp (green), spring; su (dark wheat), summer; au (red), autumn; wi (blue), winter. Lines correspond to the Poisson model for each season.

For instance, common buzzard KAI was highly significantly correlated to grassland prey index, with KAI 2.2 times higher in autumn than that in spring. In spring, during the breeding season, KAI was 4.3 times larger in the peak phase than that in the low density phase of grassland vole populations. Red kite’s correlation p-value was equal to and kestrel and hen harrier’s above but not far from the critical threshold generally accepted of p(Ho) ≤ 0.05. This lack of significance for the latter two species held from one outlier, when prey estimates were derived from the FREDON scores on a communal scale only. Dropping this observation from the data set would lead to reject Ho at p = 0.01 and p = 0.02, respectively, and to conclude formally on a correlation between the number of observations of those species and grassland prey abundance.

Fig. 6 shows the dynamics of nocturnal species. We did not detect statistically significant correlation between red fox, badger and long-eared owl abundance and grassland prey index and seasons. Domestic cat did not correlate to grassland prey index but to seasons, with lower counts in winter. Hare and wildcat KAIs were significantly correlated to grassland prey index but seasonal variations could not be detected (Table 1 and Fig. 7). Fox and hare KAIs were highly and negatively correlated to each other (p < 0.001). Furthermore, a model of hare abundance as response variable including grassland prey index and fox KAI as independent variables showed that controlling for grassland prey, hare abundance did not significantly correlate to fox KAI at a probability ≤ 0.05 (however with an observed p-value of 0.07).

**Fig 6.**
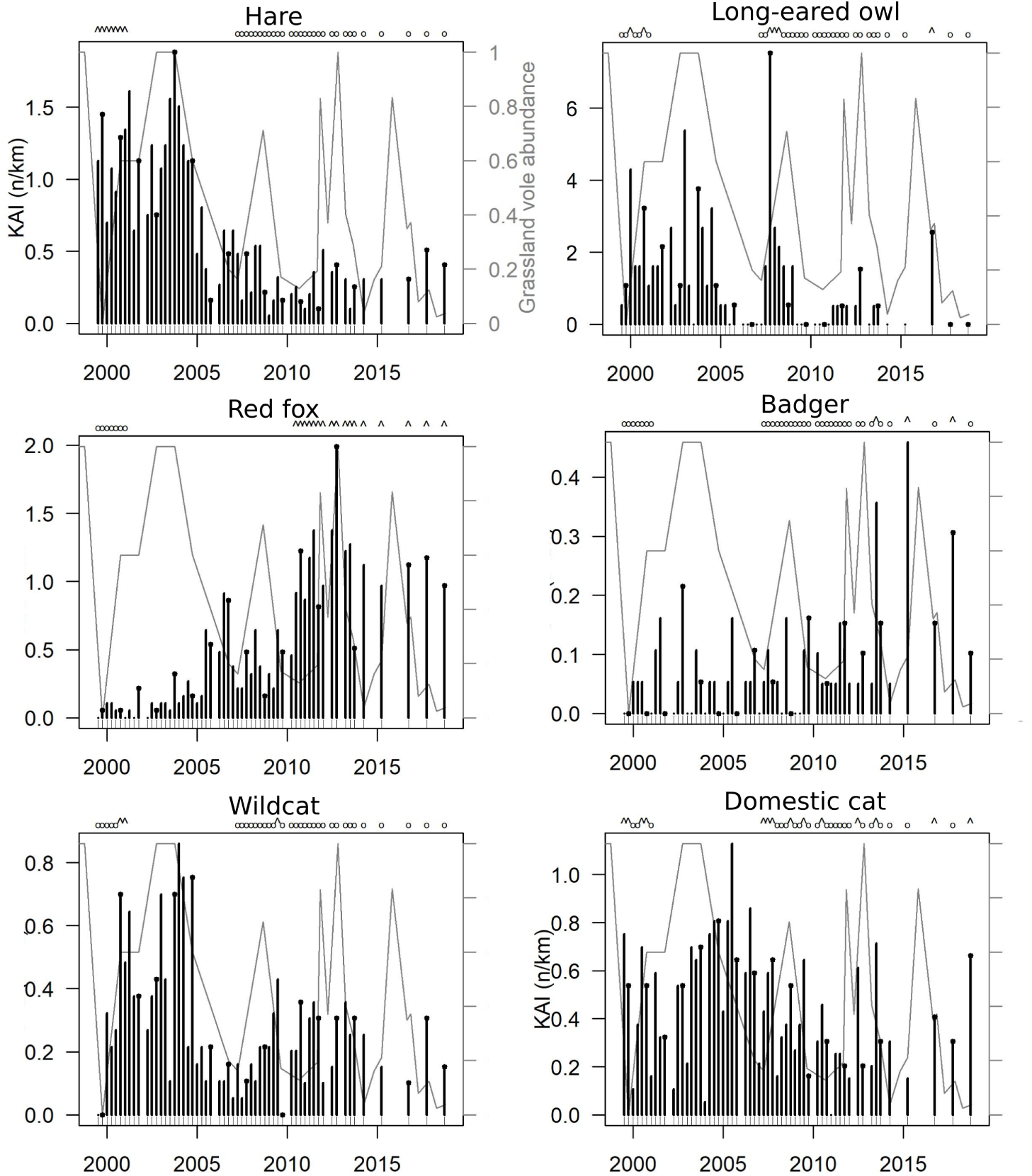
Night roadside counts. Black circles at the bar top identify autumn counts. The grey line in the background shows the variations of grassland prey abundance (the scale is the same in every plot). The letters above identify the sessions available and selected to estimate densities based on distance sampling during high (^) or low (o) abundance period.

**Fig 7.**
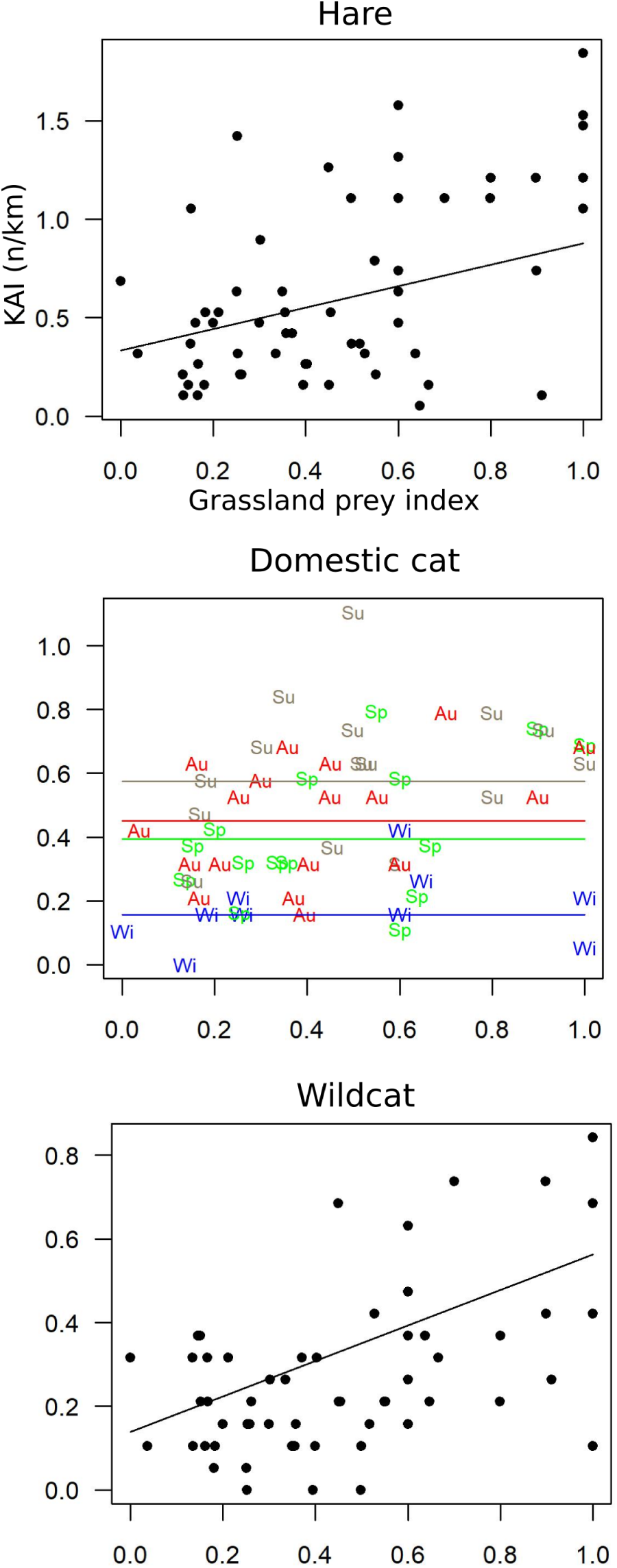
Biplots of nocturnal KAIs as a function of grassland prey index. sp (green) spring; su (dark wheat), summer; au (red), autumn; wi (blue), winter. Biplots in black have no seasonal effect. Lines correspond to the Poisson model.

Red fox and badger showed significantly higher abundance in average in the last half of the time series, and hare, wild and domestic cat, long-eared owl and hen harrier significantly lower (one-tailed permutation tests on mean, p < 0.001) (Fig. 4 and 6).

#### Spatial variations

Observations were truncated at a distance of 300 m and 350 m from the track for night and day roadside counts, respectively, accounting for 92% and 93% of their total number. Among all species in the open grassland strip along the itinerary, only the common buzzard with regard to forest and buildings, and the red fox with regard to buildings were randomly distributed. Carrion crow, red kite, kestrel and hare were observed at a greater distance to forest than expected from a random distribution; hen harrier, red fox, wildcat, long-eared owl, badger at a smaller distance; wildcat, long-eared owl and badger at a greater distance to buildings; carrion crow, red kite, kestrel, domestic cat, hen harrier at a smaller distance (Tab. 2). Seventy-five percent of the observations of domestic cat were made at less than 500 m of buildings by night and at less than 250 m by day (Fig. 8). No change in any of those patterns was observed between the first and the second half of the time series.

**Fig 8.**
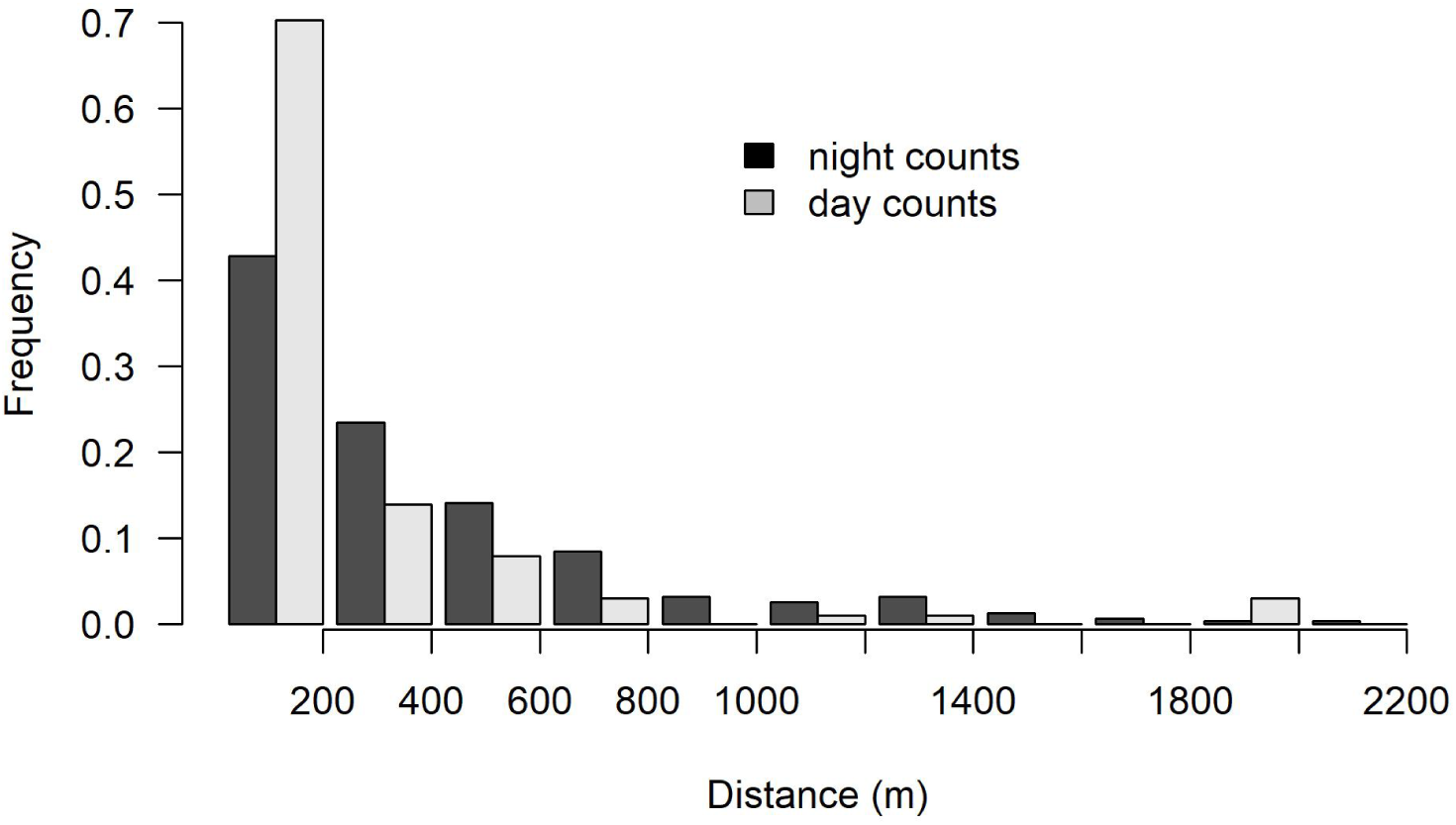
Distance to buildings of domestic cats for the night and day roadside counts (n_obs_ = 320 and n_obs_ = 101, respectively).

### Predator population density variations and daily food intake

Comparing detection models with ‘season’ as covariate with models with no covariates led us to reject the hypothesis of a seasonal effect on the detection function for every species (detection functions are presented in the annexes 1 and 2). Based on the visual examination of KAI dynamics, for each species, we identified periods when the indices could be considered similarly high or similarly low with regard to the amplitude of variations and categorize them as sub-samples of ‘low’ or ‘high’ densities (see Fig. 4 and 6). Table 3 shows conversion coefficients from KAI to densities, presents the maximum density values observed, and summarizes the estimations obtained using distance sampling by density categories (‘low’ or ‘high’). Considering the relative aggregation of the domestic cat close to buildings, we provide one density estimate for the entire study area, and another for a buffer of 300 m (night) or 250 m (day) around buildings.

**Table 2.** Mean distance (in metres) of observations to forest and buildings; random locations is the mean distance obtained from 1000 random replicates of the same number of geographical coordinates as the observations in the observation strip; the permutation test being one-tailed, p(Ho) is the number of random mean distance equal or above, or equal or below, the observed mean distance, divided by 1000. n.s., not significant. Forest effect could not be computed for the domestic cat due to its strong aggregation in and around villages

| count type | species | forest |  |  | buildings |  |  |
| --- | --- | --- | --- | --- | --- | --- | --- |
| | | observed | $p(H_0)$ | effect | observed | $p(H_0)$ | effect |
| day | carrion crow | 319 | < 0.001 | - | 852 | < 0.001 | + |
|  | common buzzard | n.s. | n.s. | = | n.s. | n.s. | = |
|  | red kite | 304 | 0.02 | - | 894 | < 0.001 | + |
|  | kestrel | 327 | < 0.001 | - | 896 | 0.002 | + |
|  | domestic cat | - | - | ? | 193 | < 0.001 | ++ |
|  | hen harrier | 236 | < 0.001 | + | 920 | 0.01 | + |
|  | random locations | 292 |  |  | 953 |  |  |
| night | hare | 299 | 0.05 | - | 874 | 0.008 | + |
|  | red fox | 253 | < 0.001 | + | n.s. | n.s. | = |
|  | domestic cat | - | - | ? | 354 | < 0.001 | ++ |
|  | wildcat | 248 | 0.002 | + | 1113 | < 0.001 | - |
|  | long-eared owl | 229 | 0.003 | + | 1043 | 0.03 | - |
|  | badger | 229 | 0.008 | + | 1125 | 0.001 | - |
|  | random locations | 284 |  |  | 939 |  |  |

**Table 3.**
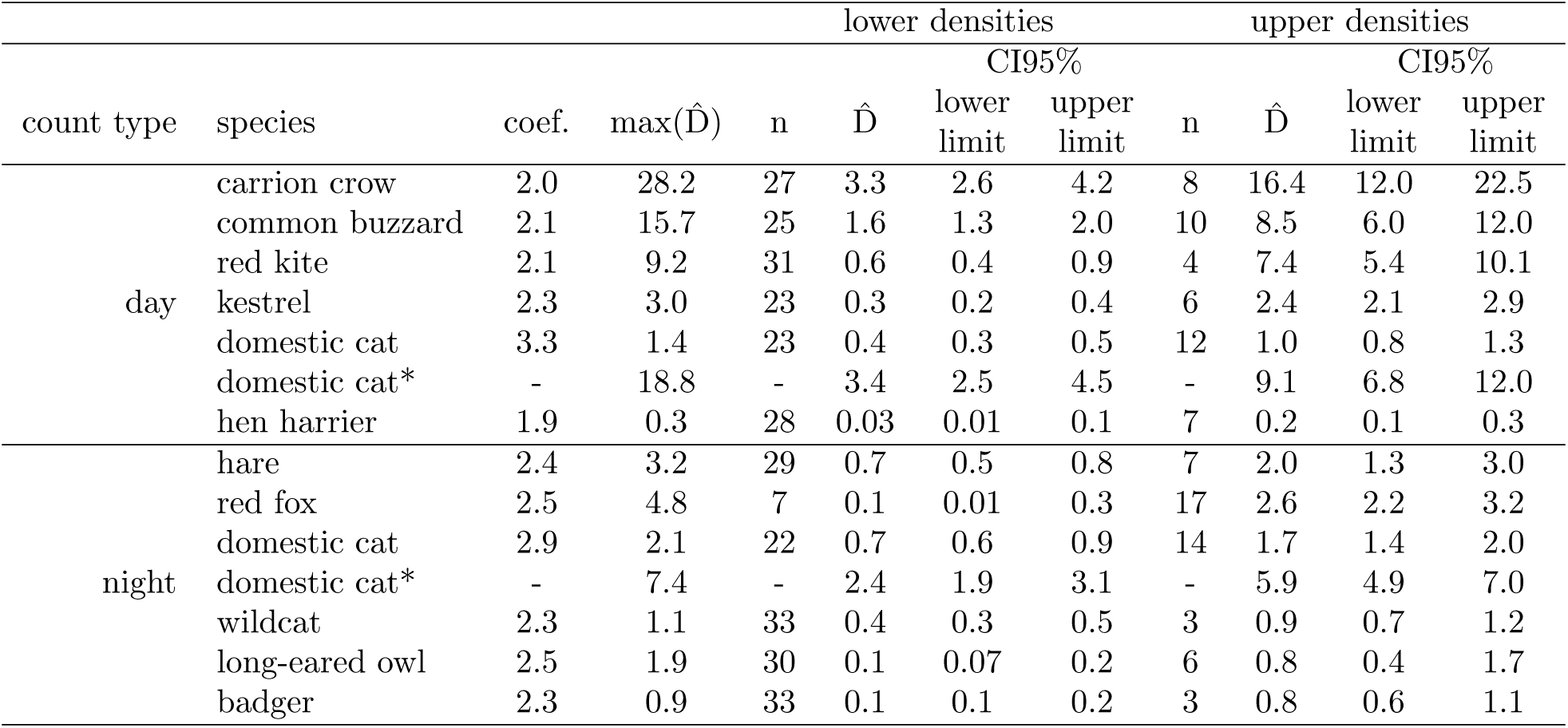
Comparison of density estimates (n.km^−2^) derived from all species data and distance sampling. Lower and upper densities correspond to estimations during low or high density period (see Fig. 4 and 6); CI95%, 95 % confidence interval; coef., conversion coefficient from KAI (n.km^−1^) into density (n.km^−2^) ; 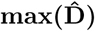, maximum density observed; n, number of sessions;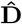, density estimate. *, domestic cat densities in a 500 m (night) or 250 m (day) buffer around buildings (including 75% of domestic cat observations, see results).

Fig. 9 shows the population density variations of the predator community during the study period for all species when both day and night road-side counts were available. Biomass and TFI variations are provided in the annexes 3 and 4.

**Fig 9.**
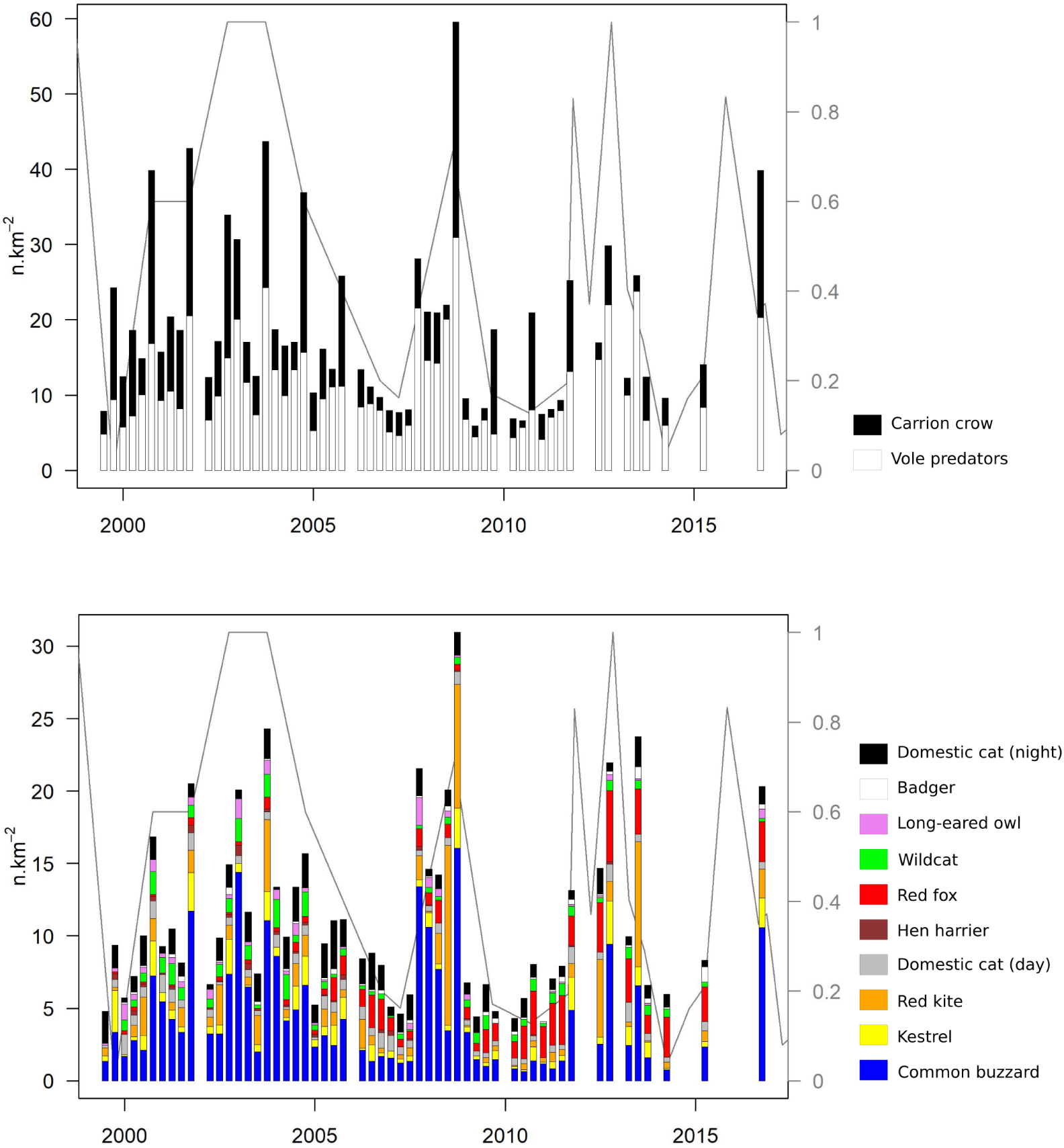
Variations in densities for each species (n.km^−2^). Variations in biomass (kg.km^−2^) and theoretical daily food intake (kg.km^−2^.day^−1^) are presented in the annexes 3 and 4.

The main features of the dynamics retained the importance of the carrion crow (range from 4.4-56.9% of the total TFI), the common buzzard (range from 4.7-48.6% of the total TFI) and the red kite (0-54.5% of the total TFI) over the entire time span, as well as the gradual increase in the red fox from 1999 to 2010 (Fig. 9). With the numerical importance of the carrion crow apart, three key periods could be identified: (i) 1999-2004 with an extremely low red fox density not exceeding 0.2 ind.km^−2^, where the community was numerically dominated by cats (domestic and wild) and common buzzards, (ii) 2005-2009 with an increasing density of foxes, and (iii) 2010-2016 with higher fox densities stabilized at an average of 2.7 ind.km^−2^. Foxes represented only 5.5% of the predator biomass (2.8% of the total TFI) in 1999-2004 but reached 29.5% (31.4% of the TFI) in 2010-2016. Regardless of the period and relative densities of species, the average TFI in the three periods was close to 4 (3.8-4.2) kg.km^−2^.day^−1^. The largest predator densities were reached during the high density peaks of grassland vole populations, with a maximum observed in autumn 2008, with 60 ind.km^−2^ (carrion crow making 48% of this total) and a daily TFI of 10.7 kg.km^−2^.day^−1^ (39.3% from carrion crow).

Table 4 summarizes the results at the grassland vole population peaks in the autumns 2003, 2008 and 2012, and in the low density phases of autumn 1999, spring 2007, autumn 2010 and spring 2014. In autumn 1999, the first 4 species totalling 91% of the TFI were the carrion crow, common buzzard, domestic cat (night) and kestrel. The common buzzard was still among those first four species in the next low density phase (spring 2007), but the proportion of TFI from birds of prey still decreased, and it was preceded by the fox, carrion crow, domestic cat and wildcat in autumn 2010 and spring 2014, with these species together making up 86% and 84% of the TFI. However, in areas with a large proportion of domestic cats roaming less than 500 m from the buildings, far from villages where domestic cats are virtually absent, fox, carrion crow and wildcat alone made up 86% of the TFI. During the first two high density phases, the carrion crow, common buzzard, red kite and domestic cat (night) made up 81 and 91% of the TFI, and in autumn 2012 during the third high density phase, the fox, common buzzard, carrion crow and domestic cat (day) alone made up 81% of the TFI. Table 4 also shows that the TFI ranged from 1.5 to 2.7 kg.km^−2^.day^−1^ in the low density phases and fromn 6.9 to 10.7 kg.km^−2^.day^−1^ in the high density peaks. Thus the TFI was multiplied by 7.1 at the maximum, while the grassland small mammal population biomass was multiplied by thousands.

**Table 4.** Density (ind.km^−2^) and theoretical daily food intake, TFI (kg.km^−2^.day^−1^) in the low (LD) and high (HD) density phases of grassland vole populations. Numbers between parentheses are percentages.

|  | species | 1999 (LD) | 2003 (HD) | 2007 (LD) | 2008 (HD) | 2010 (LD) | 2012 (HD) | 2014 (LD) |
| --- | --- | --- | --- | --- | --- | --- | --- | --- |
|  |  | autumn | autumn | spring | autumn | autumn | autumn | spring |
| density | carrion crow | 14.9 (61.4) | 19.4 (44.4) | 3.1 (39.8) | 28.6 (48) | 12.9 (61.7) | 7.9 (26.5) | 3.6 (37.6) |
|  | common buzzard | 3.3 (13.8) | 11 (25.3) | 1.2 (16) | 16.1 (27) | 1.4 (6.6) | 9.4 (31.5) | 0.7 (7.7) |
|  | kestrel | 2.9 (11.9) | 2 (4.6) | 0.3 (3.3) | 2.8 (4.6) | 1 (4.5) | 3 (10) | 0.1 (1.2) |
|  | red kite | 0.2 (0.9) | 5 (11.4) | 0.3 (4.3) | 8.5 (14.3) | 0.4 (2) | 1.4 (4.6) | 0.4 (4.4) |
|  | domestic cat (day) | 0.5 (2.2) | 0.5 (1.2) | 0.5 (6.9) | 0.9 (1.5) | 0.3 (1.6) | 1.2 (3.9) | 0.3 (3.5) |
|  | hen harrier | 0.4 (1.7) | 0.2 (0.5) | 0 (0) | 0.1 (0.2) | 0.1 (0.5) | 0.2 (0.6) | 0 (0) |
|  | red fox | 0.1 (0.5) | 0.8 (1.8) | 0.5 (6.9) | 0.4 (0.7) | 3 (14.4) | 4.9 (16.4) | 2.8 (28.9) |
|  | wildcat | 0 (0) | 1.6 (3.6) | 0.4 (4.8) | 0.5 (0.8) | 0.8 (3.9) | 0.7 (2.3) | 0.6 (6) |
|  | long-eared owl | 0.3 (1.1) | 1 (2.2) | 0 (0) | 0.1 (0.2) | 0 (0) | 0.4 (1.3) | 0 (0) |
|  | badger | 0 (0) | 0.1 (0.3) | 0.1 (1.6) | 0 (0) | 0.1 (0.6) | 0.2 (0.8) | 0.1 (1.2) |
|  | domestic cat (night) | 1.6 (6.5) | 2.1 (4.7) | 1.3 (16.5) | 1.6 (2.7) | 0.9 (4.3) | 0.6 (2) | 0.9 (9.4) |
|  | total | 24.3 | 43.7 | 7.7 | 59.5 | 20.9 | 29.9 | 9.6 |
|  | total without crow | 9.4 | 24.3 | 4.6 | 30.9 | 8.0 | 22.0 | 6.0 |
| TFI | carrion crow | 2.2 (55.1) | 2.9 (34.2) | 0.4 (23.5) | 4.2 (39.3) | 1.9 (40.9) | 1.2 (16.7) | 0.5 (18.5) |
|  | common buzzard | 0.6 (15.3) | 2 (24.1) | 0.2 (11.7) | 2.9 (27.4) | 0.3 (5.4) | 1.7 (24.7) | 0.1 (4.7) |
|  | kestrel | 0.2 (5.7) | 0.2 (1.9) | 0 (1) | 0.2 (2) | 0.1 (1.6) | 0.2 (3.4) | 0 (0.3) |
|  | red kite | 0.1 (1.3) | 1.2 (14.3) | 0.1 (4.2) | 2 (19.2) | 0.1 (2.2) | 0.3 (4.7) | 0.1 (3.5) |
|  | domestic cat (day) | 0.2 (4.8) | 0.2 (2.3) | 0.2 (10) | 0.3 (3) | 0.1 (2.6) | 0.4 (6.1) | 0.1 (4.2) |
|  | hen harrier | 0 (1.2) | 0 (0.3) | 0 (0) | 0 (0.1) | 0 (0.2) | 0 (0.3) | 0 (0) |
|  | red fox | 0.1 (1.6) | 0.4 (4.6) | 0.3 (13.3) | 0.2 (1.8) | 1.5 (31.3) | 2.4 (34) | 1.3 (46.4) |
|  | wildcat | 0 (0) | 0.6 (7.4) | 0.1 (7.5) | 0.2 (1.8) | 0.3 (6.8) | 0.3 (3.9) | 0.2 (7.9) |
|  | long-eared owl | 0 (0.6) | 0.1 (1) | 0 (0) | 0 (0.1) | 0 (0) | 0 (0.5) | 0 (0) |
|  | badger | 0 (0) | 0.1 (1.1) | 0.1 (5) | 0 (0) | 0.1 (2) | 0.2 (2.6) | 0.1 (3.2) |
|  | domestic cat (night) | 0.6 (14.3) | 0.7 (8.9) | 0.5 (23.8) | 0.6 (5.3) | 0.3 (7) | 0.2 (3.1) | 0.3 (11.3) |
|  | total | 4.0 | 8.4 | 1.9 | 10.7 | 4.6 | 6.9 | 2.9 |
|  | total without crow | 1.8 | 5.5 | 1.5 | 6.5 | 2.7 | 5.8 | 2.3 |

## Discussion

### Response of predators to grassland vole population variations

Among the 11 predator species monitored, 4 (maybe 6, if kestrel and hen harrier are included) showed a numerical response to the large variations of grassland prey observed over the 20 years of monitoring, namely, the common buzzard, red kite, wildcat and hare, as well as possibly the kestrel and hen harrier. However, responses were modulated by population trends on a larger scale. This modulation was the case for the hen harrier and long-eared owl, with populations decreasing over time in the study area reflecting the general decrease of those species in Franche-Comté and nearby Switzerland [57]. Those variations were also seasonal with generally larger populations in autumn, or in summer for the red kite, corresponding to dispersal and post-breeding migration. The numerical response of the hare, an herbivore, to the grassland vole density variations is more surprising (but see next section). A similar pattern has been observed nearby at a 30 km distance from the study area, from 1976 to 1995, for the capercaillie (*Tetrao urogallus*), in the Massif du Risoux, where the number of fledglings per hen was positively correlated to the cyclic abundance of *A. terrestris* populations [58]. This response was interpreted as being the result of predation switches during the decline phase of the voles, with a supposed relaxation of the predation pressure on the capercaillie during the high density peak, that is well documented e.g. in Scandinavian ecosystems [23, 59, 60].

The variations in the populations of other species were independent of the grassland vole populations over the study time span.

### Long term changes in the predator community structure

A striking feature of the population dynamics observed was the increase in the fox population from the beginning of the study to autumn 2010, independently of the vole populations variations. This increase can be attributed to changes in grassland small mammal control practices by farmers who shifted from late-rodenticide-only to early-integrated control in the early 2000s [44], dividing by more than forty four the quantity of anticoagulant rodenticide used during the 2010-2018 cycles compared with 1996-2000 (Fig. 3a). Massive use of anticoagulant rodenticide, here bromadiolone, is known for its deleterious side effects on vole predators [9], with a canid sensitivity that is more than 3 times higher than that of felids [61], and this effect has been proven to have drastically decreased the fox population in the area at the end of the 1990s [62] until the beginning of our study. This difference in sensitivity might explain simultaneously relatively large cat populations due to extremely limited effects of poisoning.

Furthermore, Jacquot et al. [8] have shown how the fox population has recovered on a regional scale after the change in rodent control practices. In our study, the predator community shifted from a very low fox density of 0.1 ind.km^−2^ (CI95% 0.01-0.3) foraging in grassland up to a much larger fox abundance of 2.6 ind.km^−2^ (CI95% 2.2-3.2), with a peak at 4.9 ind.km^−2^ in autumn 2012 (followed by a stabilization or a slight decrease with an epidemic of sarcoptic mange, which is still ongoing). This value is one of the highest population densities reported in rural landscapes of Europe [63, 64]. This increase was concomitant with a sudden and dramatic decrease in the hare population during a low density phase of the vole populations, and with a decrease in wild and domestic cats. This result strongly suggests that those declines might be the consequences of the increase in the fox population, possibly by direct predation or by creating a ‘landscape of fear’ [65, 66], thus limiting the distribution of the prey species to shelter-areas where they could not be detected by roadside counts (houses, forest, etc), or both. In Australia, fox removal experiments showed in one study that cats foraged more in open habitats where foxes were removed [67] and in two others that they were more abundant [68, 69]. Furthermore, in western Poland, the hare population during the same year had 1.7 times higher density in response to fox removal [70], and responded positively to sarcoptic mange epidemics that depressed the fox population in Scandinavia [23]. We did not observe changes in the spatial distribution of species between the first and second half of the study, making the ‘landscape of fear’ hypothesis less likely herein, thus suggesting a major role for direct predation.

However the long-term increase in the European badger population since the rabies vaccination in the early 1980s has been well documented in Europe [71–73]. In our study, the sudden increase since summer 2013 remains unexplained.

Excluding the stability of the carrion crow population in large numbers, a striking feature of our system is the change in predator community structure over the study period. In the early 2000s, the community was numerically dominated by the common buzzard and domestic and wildcats, and with the increase in the fox population, it became numerically dominated by the fox itself. However, foxes did not add their number to the other predators and this population increase did not lead to an increase in the average number of predators present in the study area. Large variations in vole predator number could be clearly attributed to the temporary increase in the populations of mobile birds of prey (common buzzard, red kite, etc.) in response to grassland vole outbreaks. This stability in the average predator number observed (e.g. in the low density phases of vole populations) suggests compensations among resident species due to predation or competition. Similar compensation has already been suspected in Fennoscandia, where experimental removal of avian predators to understand their role in vole population regulation led to least weasel density increase [74]. In our study, the lack of data regarding *Mustela* sp. and *Martes* sp. does not permit us to determine whether those compensations observed in a community subset extend to the whole community of vole predators. Earlier studies in the area and a nearby valley of Switzerland [75, 76] as well as in Fennoscandia [77] and northern Spain [16], clearly show that least weasel and stoat abundances follow grassland vole population peaks. Furthermore, small mustelid abundance has been shown to be dampened by fox in north America [78], by generalist predators [77] and by birds of prey [74] in Fennoscandia, but those interactions, which are possible in our study area were not unexplored herein. Moreover, small mustelids forage in vole galleries and shelter therein from bigger predators. The use of rodenticide baits buried in vole galleries as enforced by regulation [45] might contribute to an additional specific local depression of small mustelid populations [79, 80].

### Consumption of voles by predators and possible impacts on vole populations

This is the first study, to our knowledge, to provide data on the variations in population densities and daily TFI of a large community of vole predators in a temperate ecosystem in response to large variations of cyclic grassland small mammals over 20 years (four *A. terrestris* population cycles). Several biases mentioned above are inherent to the methods used; however, we consider some robust conclusions that can be carefully drawn from this exceptional long-term data set. One additional limitation derives from the observation that the functional response of each species (the dietary variations as a function of available food resources) was not studied in parallel to the variations in population densities, thus limiting the interpretation of the variations in daily TFI and the evaluation of its impact on prey populations. Thus, here we first consider first what we know about the predator diet before discussing the possible impact on the vole prey.

#### Dietary issues

The carrion crow is mostly opportunistic and feeds principally on invertebrate, cereal grain but also small vertebrates, bird eggs and carrion, in various proportions according to the place and season. At the extreme, vertebrate and eggs in particular can reach 86.6% of dry the weight of pellets in winter, e.g., in southern Spain, and they are often observed to cooperate when killing small vertebrates in pairs or small groups, also commonly forcing other birds including raptors to drop prey [38]. Their behaviour has not been systematically studied in our area, and the importance of small mammals in the diet is not yet known; however, all the behaviours mentioned above, including scavenging on dead animals, hunting voles and forcing raptors, have been occasionally observed [81]. Thus, one can hardly infer conclusions about the impact of such an opportunistic species in this ecosystem e.g., on vole regulation. Mechanically, however, their number likely has a chronic impact on species that are vulnerable to predation such as small game and bird nests.

The other species are more specialized towards small mammal prey. The detailed diet of the domestic cat is unknown in our area. However, in a similarly rural area of the Ardennes, rodents made up 55.9% of the dietary items found in 267 domestic cat faeces (6% birds, 36.7% human-linked food), with little difference between outdoor cats (owned by people other than farmers) and farm cats [82]. Rodents (Murids and Cricetids) constitute the main prey of wildcats, and they can account for 97% of the diet composition [83], while lagomorphs and birds generally appear as alternative prey. However, when the availability of lagomorphs increases, wildcats can substantially shift their diet towards them [84].

In the area, the dietary response of the red fox to variations of grassland vole relative densities differed between *M. arvalis* (no response) and *A. terrestris* (Holling’s type III-like) [85]. *M. arvalis* could make up to 60% of prey items in faeces even at very low densities (range from 0-80% of prey items over the whole range of vole densities), and *A.terrestris* showed a sigmoid increase that quickly reached a plateau (at 15% of the positive intervals of a transect -see material and methods) where it made up 40% of the dietary items on average (range from 0-80% of prey items). The description of the dietary response in this context where the two main prey abundances varied among several other alternative food resources is quite complex [86–89]. Comparisons of multi-species functional response (MSFR) models with empirical data on the red fox and barn owl showed that switching between prey depends on the proportion of the prey available among other prey (frequency dependence), as commonly thought, but also on the total amount of prey (density dependence), with a non-linear frequency and density dependent interactions [25].

#### Impact of predation on vole population abundance

In our study area, the population of the main prey species varied between 0 and approximately 1000 ind.ha^−1^ on a scale of tens of km^2^ [5, 26] and an amplitude 5-100 times larger than those observed on a similar scale in different areas worldwide [15, 77, 88, 90]. A similar amplitude has been reported locally for *M. arvalis* in alfalfa semi-permanent plots of some ha in an intensive agriculture matrix of ploughed fields of western France (50-1500 ind.ha^−1^) [91]. In our study area, two species, *A. terrestris* and *M. arvalis* had large fluctuations of similar amplitude against only one in the other systems. This ecosystem periodically offered (permanently on a large scale) an incredible biomass of several tens of kg.ha^−1^ of voles easy to access in grassland, to a large number of predator species. Here, we will attempt to understand in such system whether there are critical periods in vole population fluctuations when predation can be a key-factor controlling vole densities. At its maximum during autumn 2008, the TFI was 10.7 kg.km^−2^.day^−1^ and, hence, assuming an average weight of 80 g.vole^−1^ [43], corresponded to the equivalent of 134 montane water voles.km^−2^.day^−1^. With a carrying capacity of 1000 water voles.ha^−1^ and a predator diet made of an extreme 100% water vole at a high density of voles, exceeding 78 voles.ha^−1^, this community would not be able to decrease the vole population during its growth phase (for simulations, see https://zaaj.univ-fcomte.fr/spip.php?article114&lang=en and https://github.com/pgiraudoux/shinyPred/tree/master/shinyPred_en for the code). At densities of voles exceeding some tens of voles.ha^−1^, predators alone do not appear to be capable of instigating a population crash in our area. By contrast, daily TFI at low or medium water vole densities can substantially slow down the population increase. For instance, with a population of 2 voles.ha^−1^ at the beginning of the reproduction season, and a conservative 50% of voles in the diet for the lowest TFI (1.9 kg.km^−2^.day^−1^, hence 12 voles.km^−2^.day^−1^ in spring 2007), the model indicates that voles would be 27 ind.ha^−1^ at the end of the year instead of 91 ind.ha^−1^ without predation. The observed TFI at a low density (see 4) can even lead (based on the model) to vole extinction (e.g. autumn 1999 and 2010). Hence, to summarize, predator populations can consume several tens of thousands of voles.km^−2^.year^−1^ in our study area, and the increase in predator populations was likely not enough alone to trigger the decline in vole populations. However, our results suggest that predators during the low density phase were sufficient to considerably slow down the growth phase or even cause the extinction of vole populations locally.

Furthermore, our study documented that domestic cat populations could reach much higher densities of 2.4-9.1 ind.km^−2^ up to more than 18 ind.km^−2^ around villages within a 250-500 m radius, except during winter nights when they likely prefer to stay warmly at home. In south-central Sweden, Hansson [92] observed that domestic cats, supplied with continuous alternate food, were able to dampen the population fluctuations of the field vole, compared to more or less cat-free areas. In villages some kilometres from our study area, Delattre et al. [27, 93] reported a systematic decrease in the abundance of common vole colonies around villages near our study area during similar fluctuations of vole abundance, within an area extending 300 to 400 m from the village edge. This gradient persisted throughout a complete vole population fluctuation. They subsequently hypothesized that this lower density of voles might be the result of cat predation around villages. This figure and our estimates indicate that the combination of domestic cat density and diet, added to the density and diet of other predators, is sufficient to explain this effect.

The specific distribution of domestic cats, close to villages, can also cause spatial heterogeneity in predation pressure. For instance, during small mammal low density phases, their proportion varied between 5.9% (autumn 2010) and 23.4% (spring 2007) of the total number of predators counted.

## Conclusion

Overall, our results indicate that in such ecosystem with large variations of grassland prey, the structure of the predator community can change over the long term without changing its overall TFI variation pattern over a rodent cycle. Although the role of small and medium mustelid populations remain unknown, the higher predator densities observed during the grassland rodent peak were mostly due to mobile birds of prey that followed the rodent population increase. However, our results suggest that resident predators alone during the low density phase of grassland rodent populations were able to slow-down the increase or even to cause the extinction of rodent populations locally, but the whole predator community alone was unable to explain the population decrease observed after a high density peak. In such a system, the carrion crow was numerically the largest population with the largest TFI, but its impacts on the ecosystem could not be clearly assessed due to its eclectic diet. After a shift in rodent control practices and a much more moderate usage of anticoagulant rodenticides, the red fox population recovered and then stabilized at much larger densities, which likely negatively impacted hare, wildcat and domestic cat populations. The domestic cat population was aggregated close to buildings, with a 400 m buffer where the vole population was generally lower.

From an applied viewpoint, our results strongly suggest that, in such a highly productive and connective grassland system favourable to grassland voles, any means aimed at increasing the populations of predators during the low density phase (e.g. hedgerow networks, roosts, cats around villages, etc.) should lead to better control of grassland small mammal populations (slowing down the increase phase) [94]. However, the impacts of a management with large densities of cats around human settlements on other wildlife [95, 96] and pathogen organism transmission (e.g. *Toxoplasma gondi*) [97, 98] should be considered. Moreover, in such systems and due to unavoidable prey switches some populations such as the European hare can be caught in a predation sink and can be sustained only at low density. Management options aimed at increasing these vulnerable populations by culling predators (e.g. the red fox, etc.) would conflict with the interests of other stakeholders interested in small mammal pest control. The prohibitive costs and manpower for culling a large number of predators over the long term and the ethical concerns associated with such management should prevent this approach, which has most often been shown to be unsuccessful [99–101] and not accepted socially [102]. Other tactics should be sought, including adaptive hunting plans and demand, modification of habitats and landscapes favouring other equilibria in the community, which implies evidence-based and constructive dialogue about management targets and options between all stakeholders of such socio-ecosystems [103].

## Data accessibility

Data are available as a zip file at https://doi.org/10.1101/2020.03.25.007633. It includes:

**S1 kml file.** Location of the study area (can be dropped in a Google Earth window or read from a GIS)

**S2 Excel file.** Road side counts (sheet 1) and list of species observed (sheet 2)

**S3 Excel file.** Small mammal data

**S4 Excel file.** Data for computing theoretical daily food intakes

## Acknowledgements

This article is dedicated to Régis Defaut, who was deceased in 2008 and has been at the initiative and the true core of the ZELAC, a farmer group experimenting on environmentally-friendly control of *A. terrestris* populations in the study area, including the road side counts and transects used in this study since 1999. Special thanks are extended to Denis Truchetet and Philippe Guillemard, from the Ministry of Agriculture, who have provided support since the beginning, to Jean-Marie Curtil from the *Chambre Interdépartementale d’Agriculture Doubs - Territoire de Belfort* and Raphaël Bouquet who have provided data on the evolution of grass and milk production, to Clementine Fristch and Francis Raoul for helpful references about food intakes, to the *Maison des sciences de l’homme et de l’environnement Claude Nicolas Ledoux*, who provided technical assistance in the IGN data, and to the innumerable FREDON technicians, students and volunteers who participated to the roadside counts for their technical assistance. Legal authorization for night roadside counts was provided by the *Direction Départementale des Territoires du Doubs*. This research was carried out in the *Zone atelier Arc jurassien* https://zaaj.univ-fcomte.fr *and received financial support from the Ministry of Agriculture and from the Région de Franche-Comté*.

## Author contributions statement

PG conceived the study with Régis Defaut and designed the sampling plan and the data base. GC organized the transects and roadside counts since 2006, AL collected transect and road-side count data, managed the data base since 2014 and georeferenced the observations. PG, AL, MC and GC participated in the road-side counts. EA provided critical insights about cat ecology. PG analysed the data and wrote the MS. All authors discussed the results and reviewed the MS.

## Annexes

### 1 Detection functions of nocturnal road side counts

**Annex 1.**
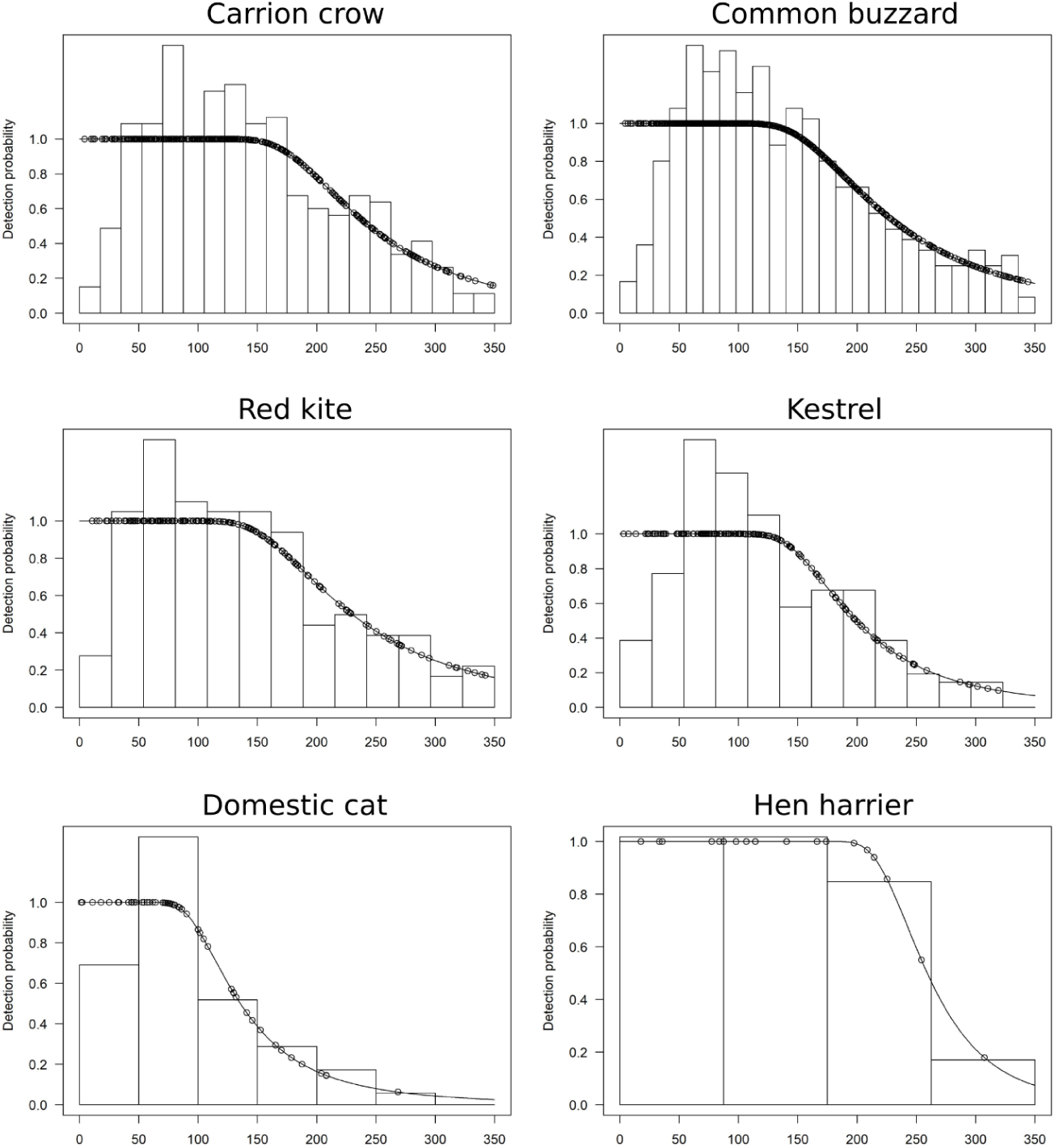
Fitted detection functions overlaid on the histogram of observed distances for diurnal species. Points indicate probability of detection for a given observation.

### 2 Detection functions of nocturnal road side counts

**Annex 2.**
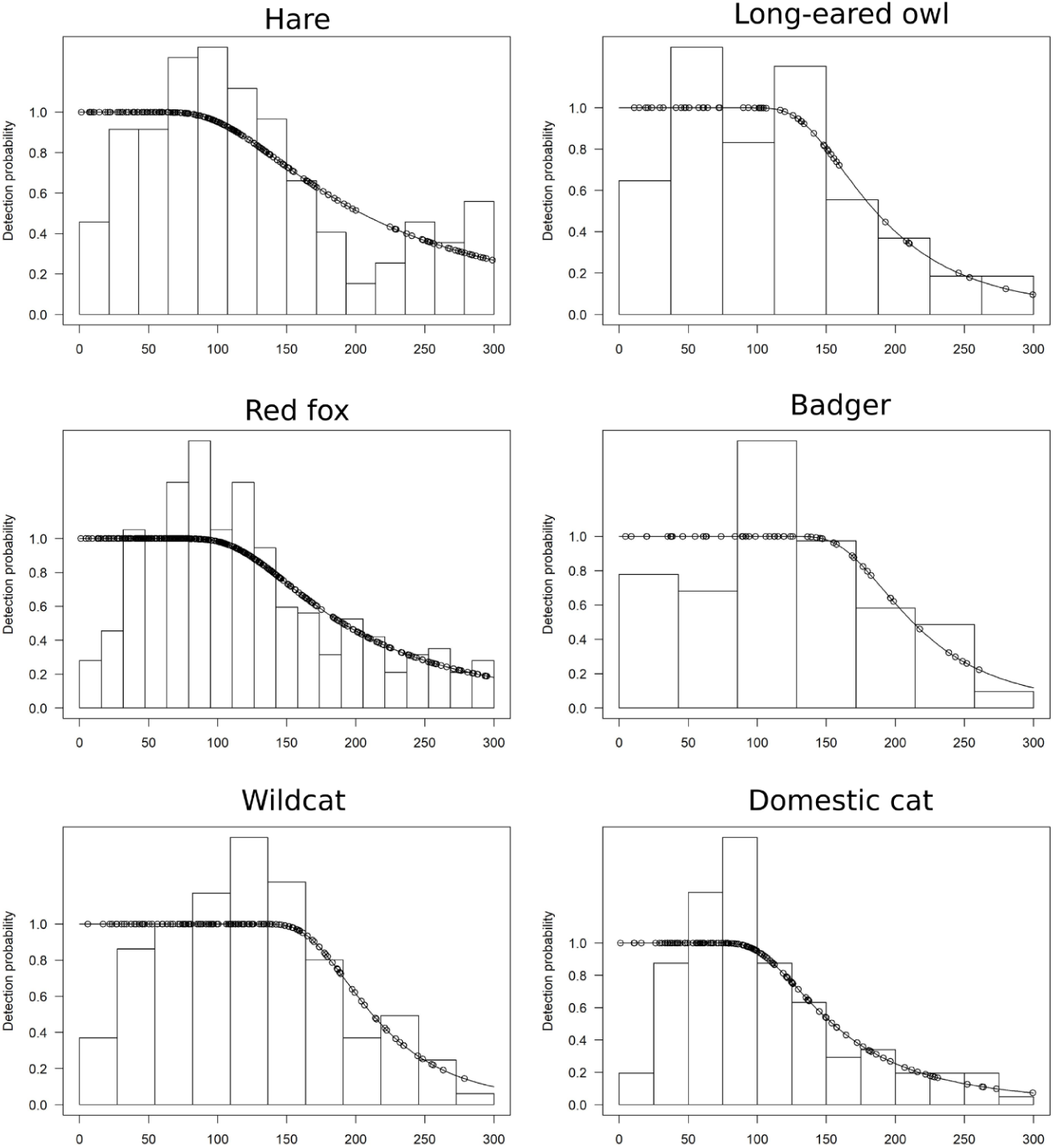
Fitted detection functions overlaid on the histogram of observed distances for nocturnal species. Points indicate probability of detection for a given observation.

### 3 Variations in biomass by species

**Annex 3.**
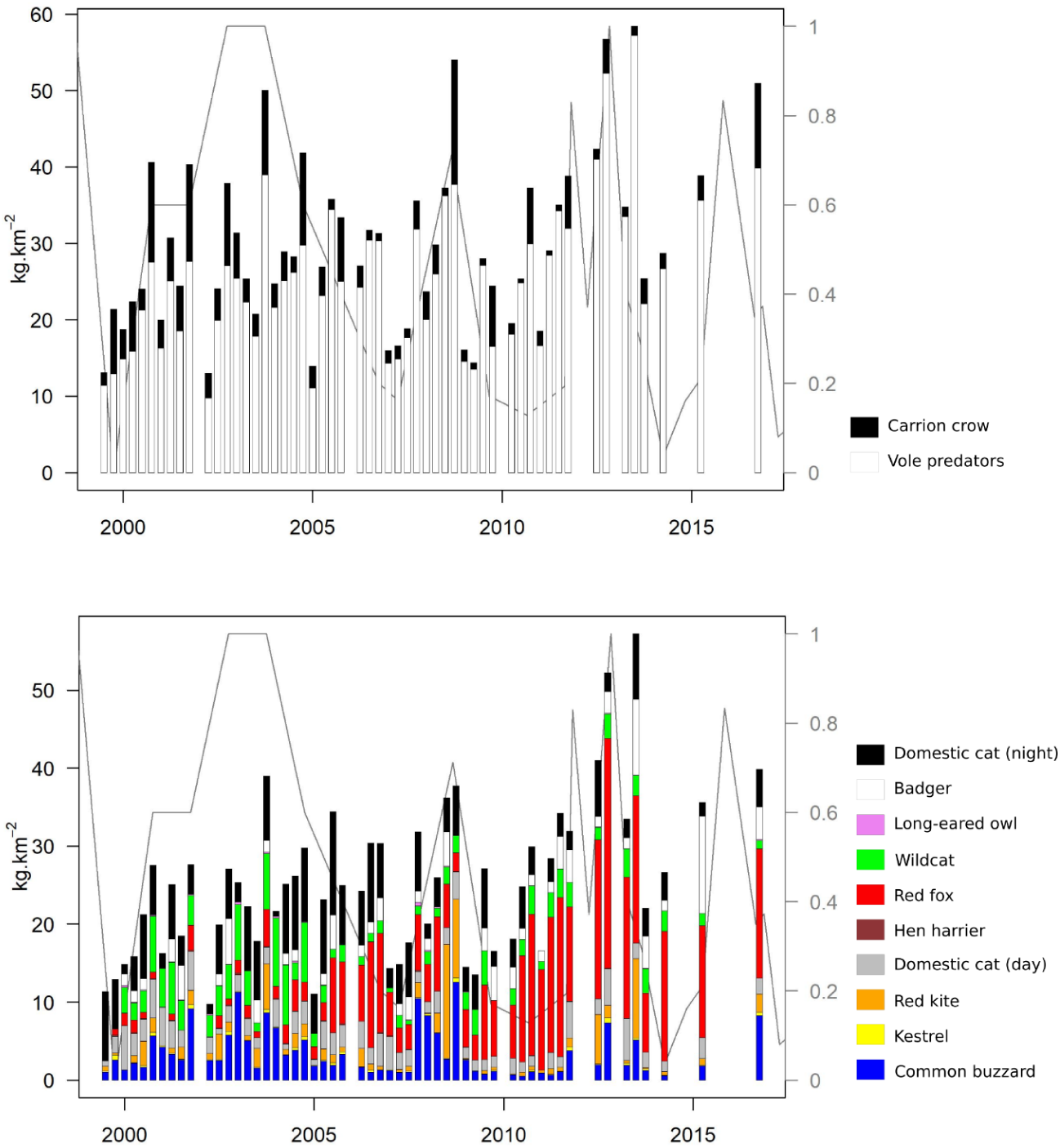

### 4 Variations in theoretical daily food intake by species

**Annex 4.**
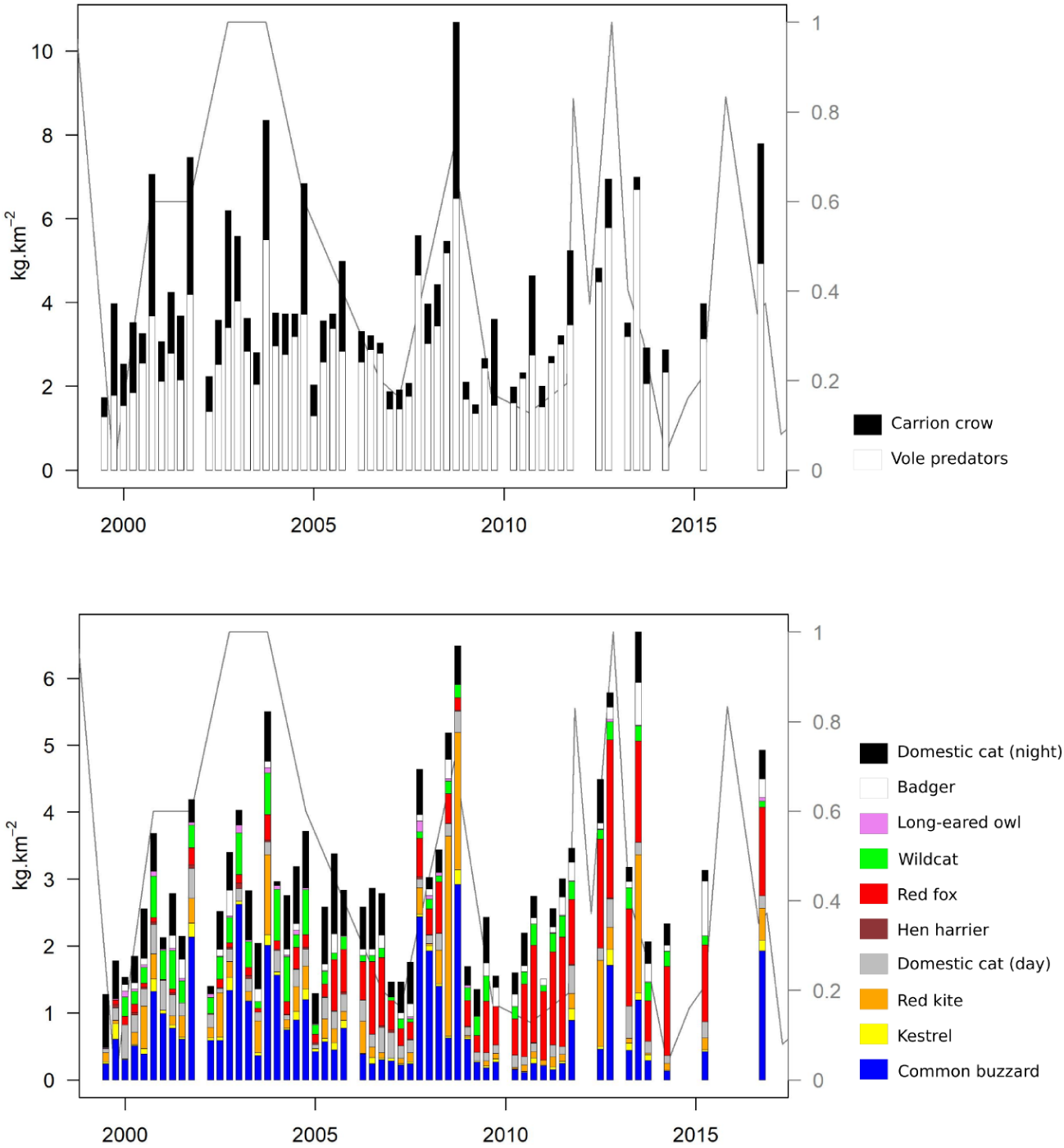

